# Loss of miRNA-153 promotes endothelial-to-mesenchymal transition and compromises lung vascular integrity

**DOI:** 10.1101/2025.10.14.682369

**Authors:** Ibrahim Elmadbouh, Zhunran Zhong, Li Wang, Michael Thompson, Luke H. Hoeppner, Y.S. Prakash, Jason X.-J. Yuan, Christina M. Pabelick, Aleksandra Babicheva

## Abstract

Endothelial-to-mesenchymal transition (EndMT) is a biological process through which lung vascular endothelial cells (ECs) transdifferentiate into mesenchymal-like cells. EndMT has recently been implicated in the development and progression of pulmonary vascular remodeling in pulmonary hypertension (PH); however, its underlying regulatory mechanisms remain incompletely understood. MicroRNAs (miRNAs) are key post-transcriptional regulators of EC gene expression and cellular responses to various stimuli. Notably, microRNA-153 (miR-153) has been shown to directly target SNAI1 to modulate epithelial-to-mesenchymal transition (EMT), a process closely related to EndMT and extensively studied in cancer. Whether miR-153 also participates in EndMT regulation, however, remains unknown. In this study, we demonstrate that 72-hour hypoxic exposure induces SNAI1-mediated EndMT in human lung vascular ECs. Hypoxia also increased cell proliferation and disrupted intercellular junctions, leading to enhanced endothelial permeability. Reduced miR-153 expression was observed in both hypoxia- and TGF-β1-induced EndMT, as well as in ECs isolated from PH patients exhibiting an EndMT phenotype. Similar to hypoxia, TGF-β1 promoted EC permeability. Loss of miR-153 enhanced SNAI1-mediated EndMT, endothelial survival, and permeability under normoxic conditions, whereas miR-153 overexpression attenuated EndMT induced by hypoxia or TGF-β1. However, miR-153 restoration did not completely recover endothelial barrier integrity disrupted by these stimuli. In conclusion, miR-153 serves as a critical regulator of EndMT, maintaining endothelial identity and barrier function. Therapeutic delivery of miR-153 may therefore represent a novel strategy to inhibit EndMT and attenuate pulmonary vascular remodeling in PH.

## Introduction

Endothelial-to-mesenchymal transition (EndMT) is a biological process in which lung vascular endothelial cells (ECs) undergo transdifferentiation into mesenchymal-like cells (1). Under physiological conditions, EndMT is a transient and tightly regulated mechanism that supports development, vascular integrity, and tissue repair. However, in pathological contexts, EndMT becomes dysregulated and contributes to the progression of several diseases, including cancer, fibrosis, and pulmonary hypertension (PH) (2-4).

EndMT-derived mesenchymal-like cells acquire a profibrotic phenotype characterized by enhanced extracellular matrix (ECM) production and remodeling activity, directly contributing to vascular fibrosis and remodeling in PH (3). The molecular profile of EndMT involves the downregulation of EC-specific markers and the upregulation of mesenchymal-specific markers. EndMT cells typically exhibit increased proliferative and migratory capacities, as well as a heightened ability to remodel the ECM.

EndMT is orchestrated by a core group of transcription factors (3, 4). Among these, the SNAI family (SNAI1/SNAIL and SNAI2/SLUG) plays a central role by repressing endothelial junctional proteins such as VE-cadherin and claudins, while activating mesenchymal genes including vimentin and fibronectin. The ZEB family (ZEB1 and ZEB2) further reinforces this transition by suppressing endothelial identity genes and promoting ECM synthesis. TWIST1 and TWIST2 serve as additional drivers, particularly under TGF-β1 or hypoxic conditions, where they regulate cytoskeletal reorganization and cell migration. These transcription factors function in overlapping and cooperative networks, often activating downstream signaling pathways such as TGF-β/SMAD, Notch, Wnt/β-catenin, and hypoxia/HIF (3). Collectively, they establish a transcriptional program that enables ECs to lose polarity and junctional integrity, acquire migratory and proliferative properties, and remodel the surrounding ECM - processes that are essential in both physiological development and pathological remodeling.

MicroRNAs (miRs) have recently emerged as critical regulators of EndMT, acting as fine-tuners of EC identity and responsiveness to profibrotic or hypoxic stimuli (1, 5). These small non-coding RNAs (∼20–24 nucleotides) modulate gene expression post-transcriptionally by binding to the 3′-untranslated region (3′-UTR) of target mRNAs, thereby inhibiting translation or promoting mRNA degradation. Several miRs have been shown to suppress EndMT in human ECs.

For example, the miR-200 family directly targets ZEB1 and ZEB2 to attenuate TGF-β1-induced EndMT in human aortic ECs (6). Similarly, miR-29 limits high-glucose-induced EndMT in human retinal microvascular ECs (7), and miR-126 preserves barrier function and maintains the phenotype of human umbilical vein ECs in response to TGF-β2 or IL-1β (8). In addition, Wang et al. demonstrated that miR-29a inhibits EndMT in an experimental model of persistent PH of the newborn (9), whereas Xu et al. reported that increased miR-126 expression promotes hypoxia-induced EndMT in a similar model (10).

Conversely, miRs such as miR-21, miR-27b, and miR-155 can promote EndMT by enhancing TGF-β signaling or repressing inhibitors of the EndMT program, particularly under hypoxic or inflammatory conditions (11). These findings highlight that miRNA-mediated modulation of EndMT in lung ECs represents a critical mechanism linking environmental and molecular cues to EC plasticity, with significant therapeutic implications for PH and fibrosis.

Among these, miR-153 has emerged as an important regulator of epithelial-to-mesenchymal transition (EMT), a process closely related to EndMT and extensively studied in cancer (12). EMT is positively correlated with cancer cell migration and invasion, leading to poor patient prognosis. Reduced miR-153 expression has been associated with advanced clinical stages and metastasis in cancer patients (13), while overexpression of miR-153 suppresses cancer cell proliferation, migration, and invasion (14, 15). Importantly, miR-153 directly targets transcription factors such as SNAI1 and ZEB2, suggesting its therapeutic potential for mitigating cancer metastasis (16). Despite these insights, the specific role of miR-153 in regulating EndMT in PH remains unknown.

The objective of this study is to determine the role of miR-153 in the development of SNAI-mediated EndMT in human lung vascular ECs.

## Methods

### Human Samples

Normal primary human lung microvascular EC were acquired as commercially available. Control and PH lung microvascular EC for comparison were provided by Pulmonary Hypertension Breakthrough Initiative (PHBI). Demographic information of human donors is provided in Table S1. Cells were cultured in T25 flasks pre-coated with 0.1% gelatin and maintained in VascuLife^®^ basal medium without antimicrobials or phenol red, supplemented with the VascuLife^®^ VEGF-Mv LifeFactors Kit (Lifeline Cell Technologies). Cells were maintained at 37°C in a humidified atmosphere with 5% CO and used at passages 3–6 for all experiments. For treatments, EC were exposed to either normoxic (21% O) or hypoxic (3% O) conditions for 72 hours using C-chamber (BioSpherix) and ProOx P110 Oxygen Single Chamber Controller (BioSpherix). In another set of experiments cells were treated with vehicle or TGF-β1 (10 ng/mL; R&D Systems) for 7 days to induce EndMT as published before (17).

### Immunoblotting

Proteins samples were extracted using RIPA buffer mixed with 6×SDS loading buffer (Boston BioProducts) and denatured at 100°C for 7 minutes (17). Protein samples were separated on 12% Bis-Tris Mini-PROTEAN TGX precast gels (Bio-Rad) and transferred onto PVDF membranes (0.45 µm; Thermo Scientific) at 96V for 2 hours. Membranes were blocked in 5% non-fat milk in TBS-T (Tris-buffered saline with 0.1% Tween 20) for 1 hour and incubated overnight at 4°C with primary antibodies: anti-SNAI1, anti-SNAI2, anti-PECAM-1, anti-CDH5, anti-VIMENTIN, anti-TAGLN, anti-PCNA (Cell Signaling, 1:1000), or β-actin (Santa Cruz Biotechnology, 1:1000). After washing, membranes were incubated with HRP-conjugated secondary antibodies (anti-mouse or anti-rabbit, 1:2500; Cell Signaling) in 2.5% non-fat milk for 1 hour. Detection was performed using SuperSignal™ West Femto Substrate (ThermoFisher) and visualized using a ChemiDoc MP imaging system (Bio-Rad). Band intensities were quantified using ImageJ software.

### Quantitative PCR (qPCR)

Total RNA was extracted using QIAzol reagent (Invitrogen) and quantified with a NanoDrop 1000 spectrophotometer (Thermo Scientific) (17). cDNA samples were synthesized using the miRCURY LNA™ RT Kit (Qiagen), and miRNA expression analysis was performed with miRCURY LNA™ SYBR Green PCR Kit (Qiagen) in a CFX384 Touch Real-Time PCR System (Bio-Rad). Primers for hsa-miR-153-3p and U6 (endogenous control) were purchased from Qiagen. The amplification protocol included: 25°C for 30 min, 42°C for 30 min, and 85°C for 5 min. The relative expression levels were calculated as described before (18).

### In Silico Analysis

*In silico* analysis was performed using the standard online software TargetScanHuman v8.0 (http://www.targetscan.org/) to predict the potential miRNAs binding the targeted mRNA as published before (18).

### Cell Transfection

Normal cells were transfected with miRCURY LNA™ hsa-miR-153-3p inhibitor or mimic (Qiagen) for 6 hours as published before (18). All-Stars Control siRNA (Qiagen) and miRCURY LNA™ miRNA inhibitor negative control (Qiagen) was used as a negative control (NC) for miRNA mimic and miRNA inhibitor transfection experiments, respectively. Following transfection, the medium was replaced, and cells were treated to induce hypoxia- or TGF-β1-induced EndMT as described above.

### Cell Viability and Proliferation Assays

Cell viability was assessed using MTS assay kit (Abcam) according to the manufacturer’s instructions. Optical density (OD) was measured at 490 nm using a Synergy Neo2 multimode microplate reader (BioTek Instruments). Data were normalized to control wells. Cell proliferation was evaluated using the Click-iT™ EdU Cell Proliferation Assay Kit (EdU-594, Invitrogen, Millipore Corp, USA) as we published before (19). EdU-positive nuclei were visualized using a Zeiss Axio Observer widefield fluorescence microscope. Images were acquired using a 20× objective and analyzed using ImageJ software. Results are expressed as the percentage of EdU-positive cells normalized to the total cell number.

### Cell Apoptosis Assay

Apoptotic cells were detected using the TUNEL assay (In Situ Cell Death Detection Kit, Fluorescein, Roche) as we published before (20). TUNEL-positive cells were imaged using a Zeiss Axio Observer microscope at 20× magnification and analyzed using ImageJ software. Data is presented as the percentage of TUNEL-positive cells normalized to the total cell number.

### Endothelial Permeability Assay

For the permeability assessment lung vascular EC were seeded on 0.1% gelatin-coated Transwell inserts (6.5 mm diameter, 0.4 μm pore size; Corning Inc.). The diffusion of FITC-dextran (70 kDa, 1 mg/mL; Sigma-Aldrich) across the EC monolayer was measured after 1 hour of incubation (21). Media from the lower chamber were transferred to black 96-well plates (Invitrogen) and fluorescence was measured at 485 nm (excitation) and 535 nm (emission) using a Synergy Neo2 plate reader (BioTek Instruments). Reference standards of FITC-dextran were included for calibration.

### Immunocytochemistry

Lung vascular EC were cultured on 2-chamber glass slides, transfected and/or treated as described above (17). Cells were fixed with 4% paraformaldehyde in PBS for 20 minutes at room temperature, followed by blocking with 3% hydrogen peroxide in methanol for 10 minutes. After washing, nonspecific binding was blocked with 10% goat serum in PBS for 1 hour at room temperature. Cells were then incubated overnight at 4°C with primary antibodies (mouse anti-SNAI1, rabbit anti-SNAI2, rabbit anti-ACTA2, or mouse anti-PECAM-1) diluted in 10% goat serum. The following day, cells were incubated with Alexa Fluor™ 488 or 568-conjugated secondary antibodies (ThermoFisher) for 1 hour at room temperature. Coverslips were mounted with DAPI-containing mounting medium (Vector Laboratories) and images were acquired using a Zeiss Axio Observer microscope with an Apotome camera.

### Statistical Analysis

All data are presented as mean ± standard deviation (SD). Statistical significance between two groups was assessed using paired or unpaired Student’s *t*-tests. For comparisons among three or more groups, one-way ANOVA followed by appropriate post hoc tests was used. Differences were considered statistically significant at *P* ≤ 0.05.

## RESULTS

### Hypoxia induces EndMT and decreases miR-153 in human lung vascular EC

Cells undergoing EndMT are losing EC-specific markers and function while gaining mesenchymal markers and the increased proliferative index. To investigate whether hypoxia stimulates EndMT in human EC we compared EndMT profile (transcription factors SNAI1 and SNAI2 as well as EC specific markers and mesenchymal markers) in normoxic and hypoxic cells. As shown in Figure 1A and B, the protein levels of SNAI1 and SNAI2 were significantly increased in hypoxic cells (1.4±0.08, p=0.023 and 2.0±0.20, p=0.003, respectively) compared with controls. Concurrently, hypoxic exposure resulted in the downregulation of EC-specific markers (PECAM-1 and CDH5) and the upregulation of mesenchymal markers, smooth muscle cell-specific marker TAGLN and fibroblast-specific marker VIMENTIN. These data demonstrate that hypoxia promotes EndMT in human lung vascular EC.

**Figure 1.**
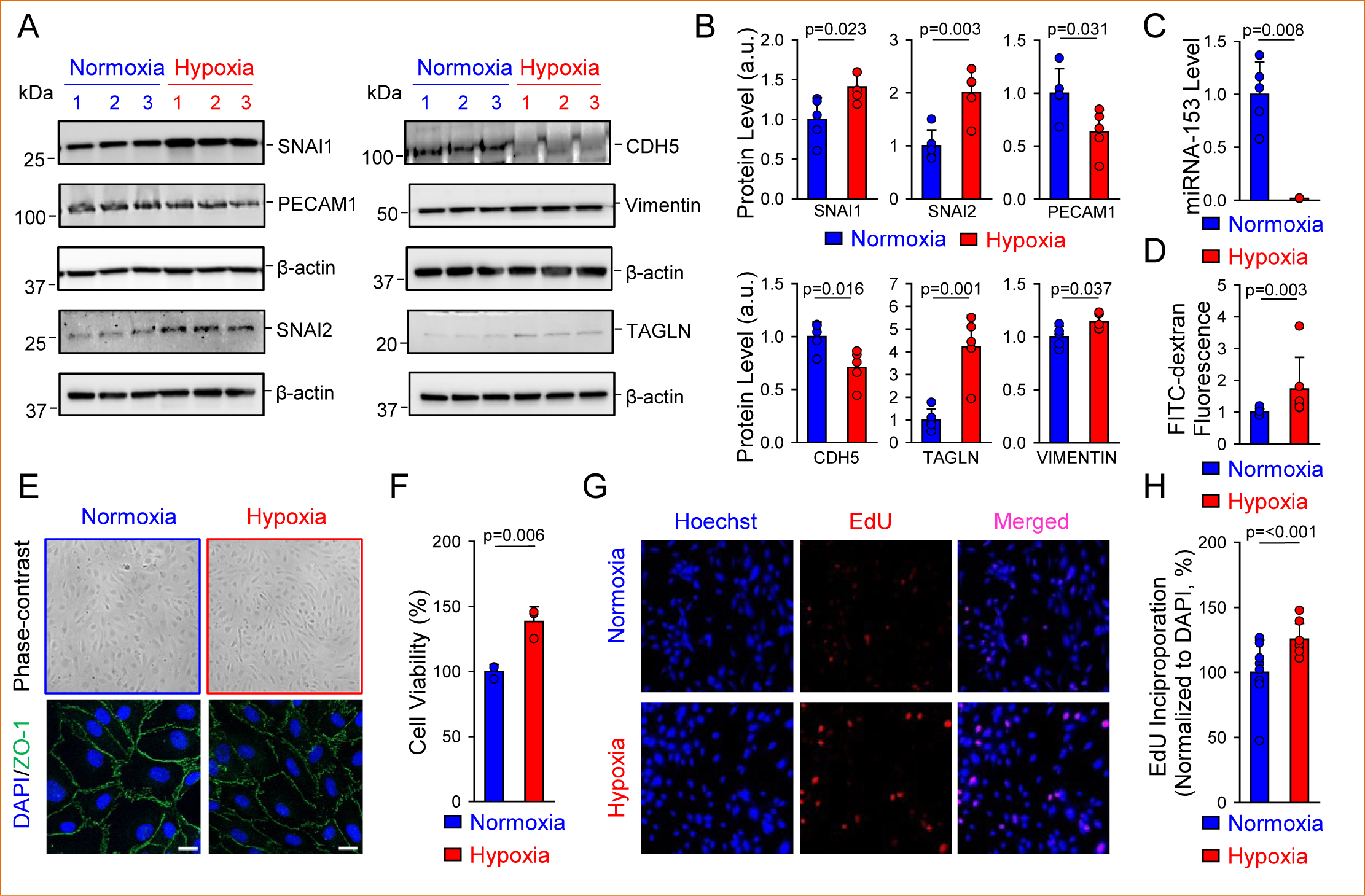
Hypoxia promotes EndMT, cell survival, and permeability while decreasing miR-153. **A-B:** Representative Western blot images (A) and summarized data (B) showing protein expression of SNAI1, SNAI2, PECAM1, CDH5, TAGLN, and VIMENTIN (normalized to β-actin) in human lung vascular EC exposed to normoxia (21% O_2_) or hypoxia (3% O) for 72 hours. **C:** Summarized data showing RT-PCR analysis of miR-153 expression (normalized to U6) in normoxic or hypoxic EC. **D:** Summarized data showing mean fluorescence intensity for FITC-dextran diffused through the endothelial monolayer of normoxic or hypoxic EC. **E:** Representative phase-contrast (top) and immunofluorescence (bottom) images showing cell morphology and ZO-1 staining respectively in normoxic or hypoxic cells. Scale bar is 20 μm. **F:** Summarized data showing cell viability using MTS assay in normoxic or hypoxic EC. **G-H:** Representative images (G) and summarized data showing EdU incorporation (H) in normoxic or hypoxic EC. Values are mean±SD (n=5 per group).

To determine whether miR-153 regulates EndMT, we used qPCR experiments to compare miR-153 expressions in normoxic and hypoxic EC. Indeed, hypoxia significantly reduced miR-153 by 98.4±0.02% (Fig. 1C). We then performed *in silico* analysis to determine the binding site for miR-153 within 3’-UTR of mRNA encoding SNAI. *In silico* analysis showed that miR-153 had the most favorable score to target 3’-UTR of SNAI1 mRNA (-0.49 or 99 percentile) (Table S2) while SNAI2 had no binding sites specific for miR-153 (Table S3). Thus, enhanced SNAI-mediated EndMT is correlated with decreased miR-153 level in hypoxic EC.

### Hypoxia stimulates endothelial permeability, survival, and proliferation

Endothelial dysfunction is considered as an initial trigger in the development of PH. To evaluate the hypoxic effect on endothelial function, we assessed barrier function, proliferation and survival rates in lung vascular EC exposed to normoxic or hypoxic conditions. As shown in Fig. 1D, the monolayer of hypoxic EC had significantly higher permeability to FITC-dextran compared with normoxic cells. We observed remarkable morphological changes in EC cultured under hypoxia (Fig. 1E). Under normoxia, EC displays the characteristic cobblestone-like morphology with uniform cell-cell contacts and minimal intercellular gaps. In contrast, hypoxia induces elongation toward the spindle-shaped phenotype, accompanied by a loss of the regular monolayer pattern and increased heterogeneity in cell orientation (Fig. 1E, *top panel*). Moreover, immunostaining experiments demonstrated tight ZO-1 organization in normoxic cells and disrupted ZO-1-positive cell-cell junctional structures in hypoxic cells indicating tight junction instability (Fig. 1E, *bottom panel*). These morphological changes are consistent with cytoskeletal reorganization and EndMT. In addition, 72 hours of hypoxic exposure elevated cell viability (Fig. 1F) and proliferation (Fig. 1G and H) suggesting enhanced survival rates. Specifically, our data indicates that hypoxia shifts EC from a quiescent, homeostatic state toward an activated, maladaptive phenotype with apoptosis resistance, excessive proliferation, and compromised barrier integrity.

### Loss of miR-153 promotes EndMT, cell survival, and barrier instability

To determine the role of miR-153 in regulating SNAI-mediated EndMT, we applied a loss- or gain-of-function approach in human lung vascular EC. At first, we assessed EndMT profile in cells transfected with NC inhibitor or miR-153 inhibitor and exposed to normoxic conditions for 72 hours. As shown in Fig. 2A and B, immunocytochemistry experiments revealed that miR-153 deficiency led to the significantly increased EndMT profile characterized by the upregulation of SNAI1 (4.19±0.11, *p*=<0.001), SNAI2 (1.37±0.06, *p*=<0.001), ACTA2 (3.24±0.23, *p*=<0.001) and downregulation of PECAM-1 (0.57±0.02, p=<0.001). In normal EC PECAM-1 staining is localized to adherens junctions that form continuous cell-cell contacts. EC transfected with NC inhibitor exhibited continuous, linear adherens junctions along cell-cell borders as visualized by PECAM-1 immunostaining (Fig. 2C). In contrast, hypoxia induced a marked disruption of junctional continuity, with PECAM-1 staining appearing discontinuous and fragmented, consistent with junctional destabilization and increased intercellular gaps. Thus, EC transfected with miR-153 inhibitor failed to form continuous adherens junctions compared with negative controls (Fig. 2D). Next, we evaluated barrier function in both groups to demonstrate that cell monolayer permeability was also significantly increased in cells with miR-153 inhibition (Fig. 2E). Transfection efficiency was confirmed by qPCR experiments. The level of miR-153 was significantly reduced in miR-153 inhibitor-transfected cells compared with negative controls (Fig. 2F). These results showed that loss of miR-153 was sufficient to induce EndMT in human EC under normoxic conditions.

**Figure 2.**
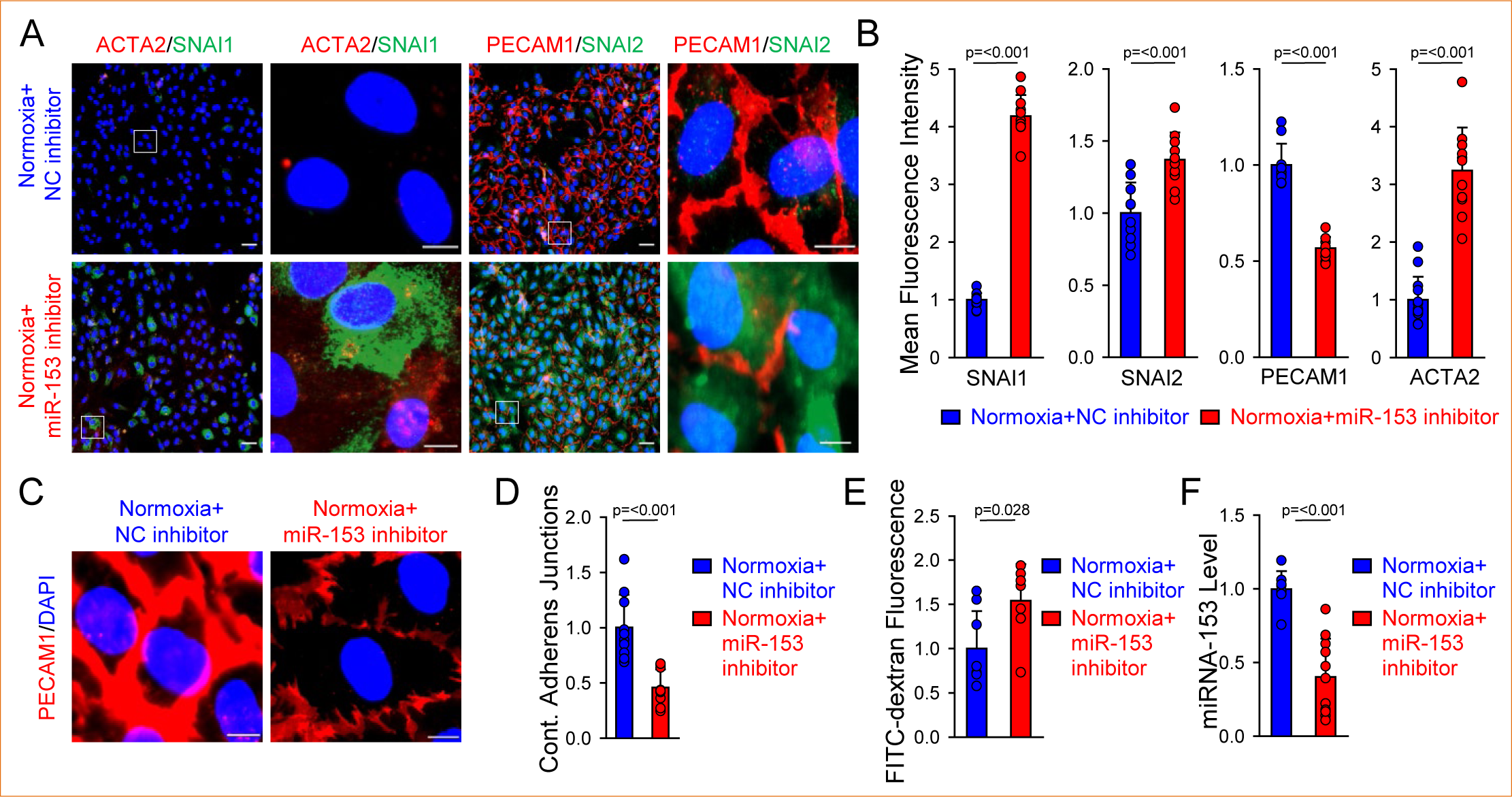
Loss of miR-153 stimulates EndMT and compromises barrier integrity. **A-B:** Representative images (A) and summarized data (B) showing mean fluorescence intensity for SNAI1, SNAI2, PECAM1, and ACTA2 in EC transfected with negative control (NC) or miR-153 inhibitor and exposed to normoxia for 72 hours (n=11-13 per group). Nuclei were counterstained with DAPI. Scale bars: 50 µm and 10 µm. **C-D:** Representative images (C) and summarized data (D) showing immunofluorescence staining for PECAM1 and the fraction of continuous adherens junctions in EC transfected with NC or miR-153 inhibitor and exposed to normoxia for 72 hours (n=9-10 per group). Scale bar: 10 µm. **E:** Summarized data showing mean fluorescence intensity for FITC-dextran diffused through EC monolayer transfected with NC or miR-153 inhibitor and exposed to normoxia for 72 hours (n=7-8 per group). **F:** Summarized data showing RT-PCR analysis of miR-153 expression (normalized to U6) in EC transfected with NC or miR-153 inhibitor and exposed to normoxia for 72 hours (n=8-11 per group). Values are mean±SD.

To determine if miR-153 deficiency affects cell survival, we compared DNA synthesis and DNA fragmentation in EC transfected with NC or miR-153 inhibitor and cultured in the medium supplemented with 2% FBS. EdU incorporation in EC with miR-153 loss was 2.77-fold greater than that in control cells indicating higher cell proliferation (Fig. 3A and B). TUNNEL assay detected lower cleaved genomic DNA in cells transfected with miR-153 inhibitor suggesting reduced apoptosis (Fig. 3C and D). The enhanced proliferation rate of miR-153 inhibitor-transfected EC was also associated with upregulated protein expression level of PCNA (Fig. 3E and F). The findings led us to conclude that miR-153 deficiency contributes to the conversion of normal slowly growing fully differentiated EC into highly proliferative apoptosis-resistant mesenchymal cells.

**Figure 3.**
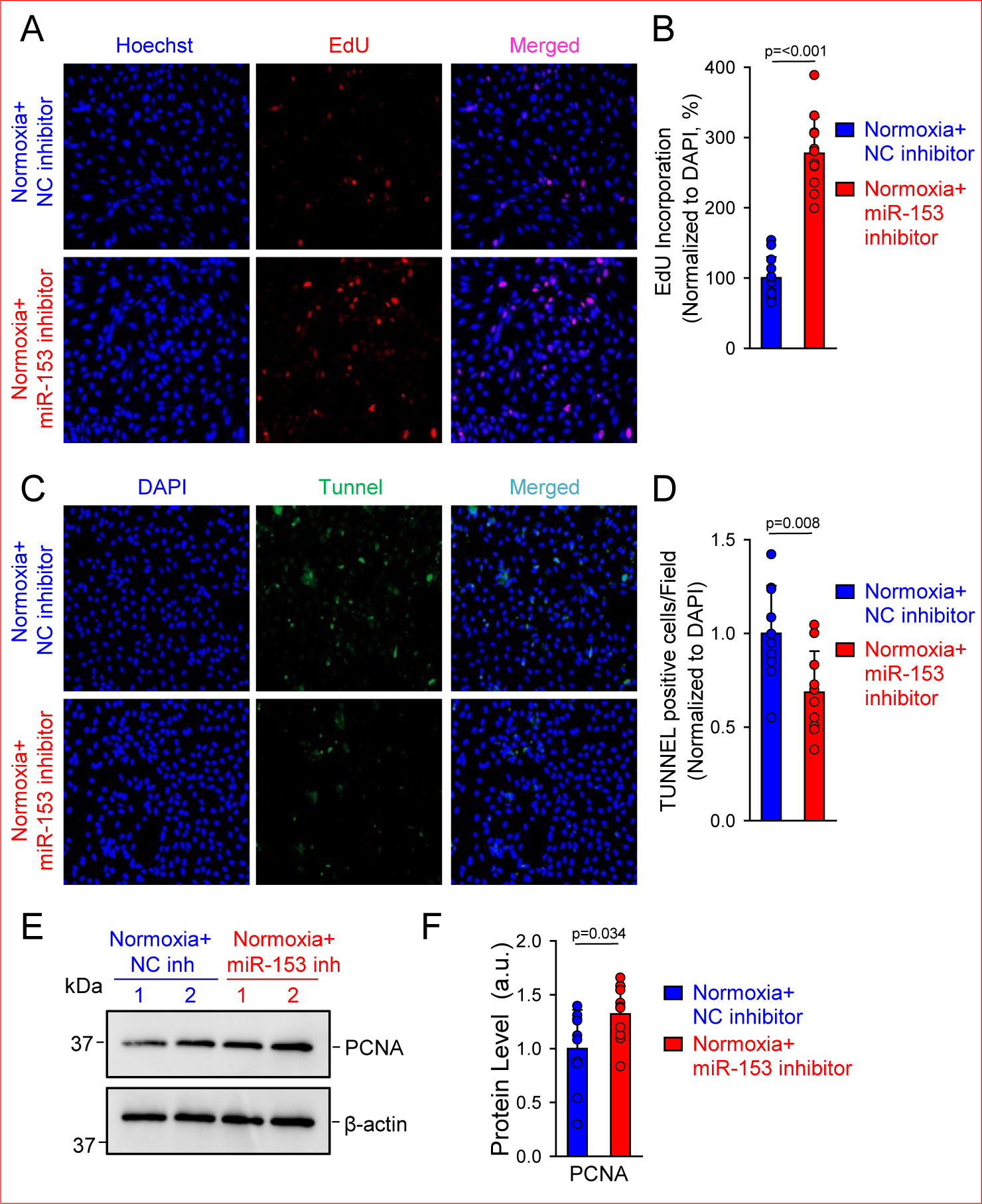
Inhibition of miR-153 enhances cell survival. **A-B:** Representative images (A) and summarized data (B) showing EdU incorporation in EC transfected with NC or miR-153 inhibitor and exposed to normoxia for 72 hours (n=12 per group). **C-D:** Representative images (C) and summarized data (D) showing TUNEL staining in EC transfected with NC or miR-153 inhibitor and exposed to normoxia for 72 hours (n=10 per group). **E-F:** Representative Western blot images (E) and summarized data (F) showing PCNA protein expression (normalized to β-actin) in EC transfected with NC or miR-153 inhibitor and exposed to normoxia for 72 hours (n=10 per group). Values are mean±SD.

### Overexpression of miR-153 attenuates hypoxia-induced EndMT, cell survival but has no effect on barrier instability

This set of experiments was designed to determine the potential of miR-153 mimic to improve hypoxia-induced endothelial dysfunction. As shown in Fig. 4A and B, transfection of EC with miR-153 mimic followed by hypoxic exposure for 72 hours significantly suppressed the protein expression of SNAI1 (0.09±0.009, *p*=<0.001), SNAI2 (0.75±0.03, *p=*<0.001), and ACTA2 (0.15±0.03, *p=*<0.001) while restored PECAM-1 level (1.43±0.03, *p=*<0.001). However, miR-153 was not able to improve hypoxia-induced disruption of junctional continuity (Fig. 4C and D). PECAM-1 staining was discontinuous and fragmented in hypoxic cells overexpressed for miR-153 indicating that miR-153 mimic failed to stabilize adherens junctions and decrease intercellular gaps. Moreover, permeability to FITC-dextran was not significantly different in cells with or without miR-153 overexpression (Fig. 4E). The transfection efficiency was confirmed by qPCR experiments. The miR-153 expression was dramatically elevated in cells transfected with miR-153 mimic compared with NC controls (Fig. 4F). These results demonstrated that miR-153 was able to abolish hypoxia-induced EndMT but was not sufficient to improve hypoxia-disrupted endothelial barrier integrity.

**Figure 4.**
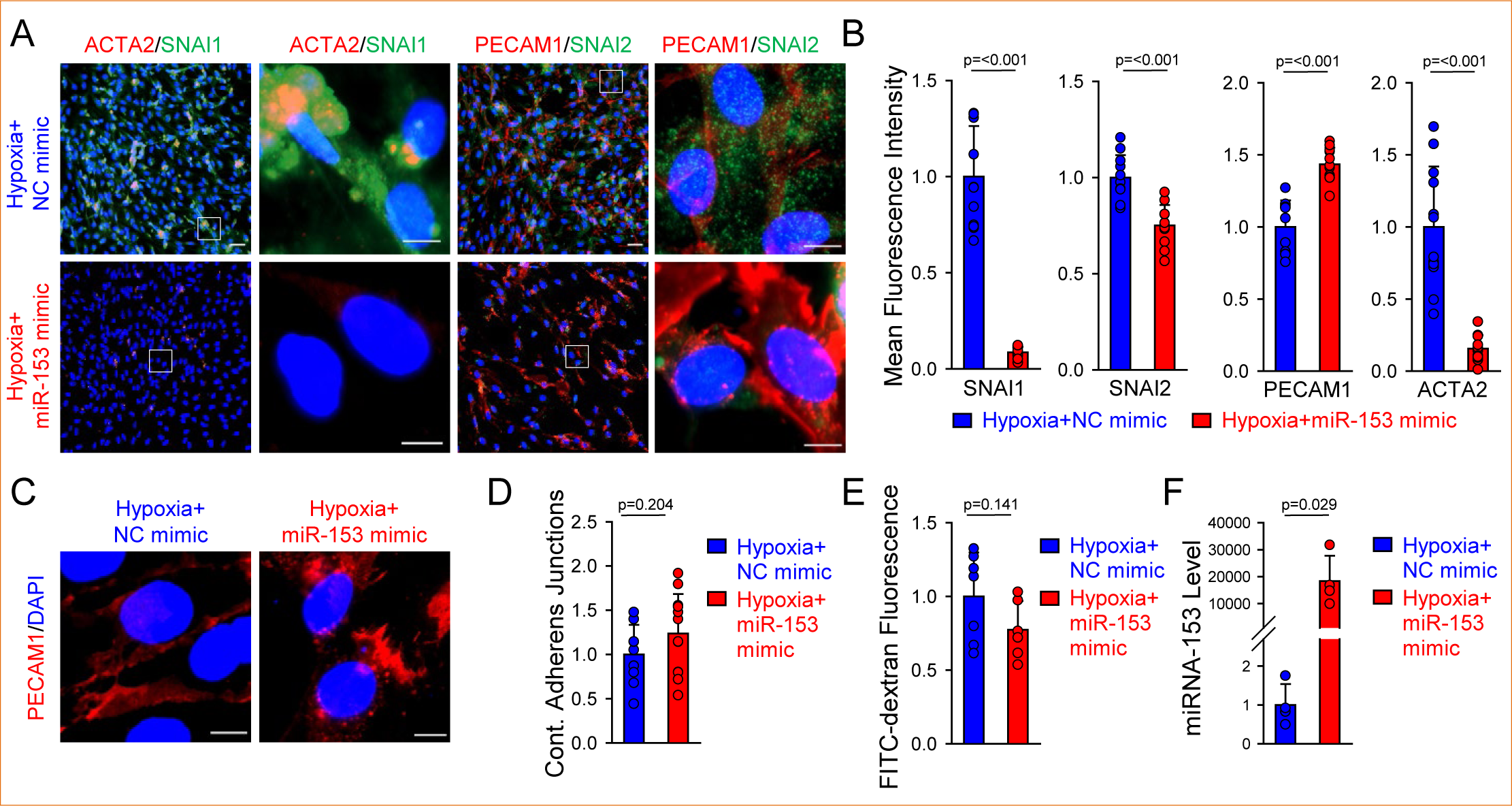
miR-153 attenuates hypoxia-induced EndMT and cell permeability. **A-B:** Representative images (A) and summarized data (B) showing mean fluorescence intensity for SNAI1, SNAI2, PECAM1, and ACTA2 in EC transfected with NC or miR-153 mimic and exposed to hypoxia (3% O) for 72 hours (n=11-12 per group). Nuclei were counterstained with DAPI. Scale bars: 50 µm and 10 µm. **C-D:** Representative images (C) and summarized data (D) showing immunofluorescence staining for PECAM1 and the fraction of continuous adherens junctions in EC transfected with NC or miR-153 mimic and exposed to hypoxia for 72 hours (n=9-12 per group). **E:** Summarized data showing mean fluorescence intensity for FITC-dextran diffused through EC monolayer transfected with NC or miR-153 mimic and exposed to hypoxia for 72 hours (n=6-7 per group). **F:** Summarized data showing RT-PCR analysis of miR-153 expression (normalized to U6) in EC transfected with NC or miR-153 mimic and exposed to hypoxia for 72 hours (n=4 per group). Values are mean±SD.

Next, we examined whether miR-153 prevented hypoxia-induced survival in human lung vascular EC. The experiments using EdU assay showed the reduced cell proliferation by 40% in cells transfected with miR-153 mimic compared with NC-transfected cells (Fig. 5A and B), while the experiments with TUNEL assay revealed significantly increased apoptosis (Fig. 5C and D). Also, EC overexpressed for miR-153 exhibit lower protein expression of PCNA compared with controls (0.71±0.09, *p* = 0.048, Fig. 5E and F). Overall, these findings suggest that miR-153 abolished endothelial dysfunction in response to hypoxia.

**Figure 5.**
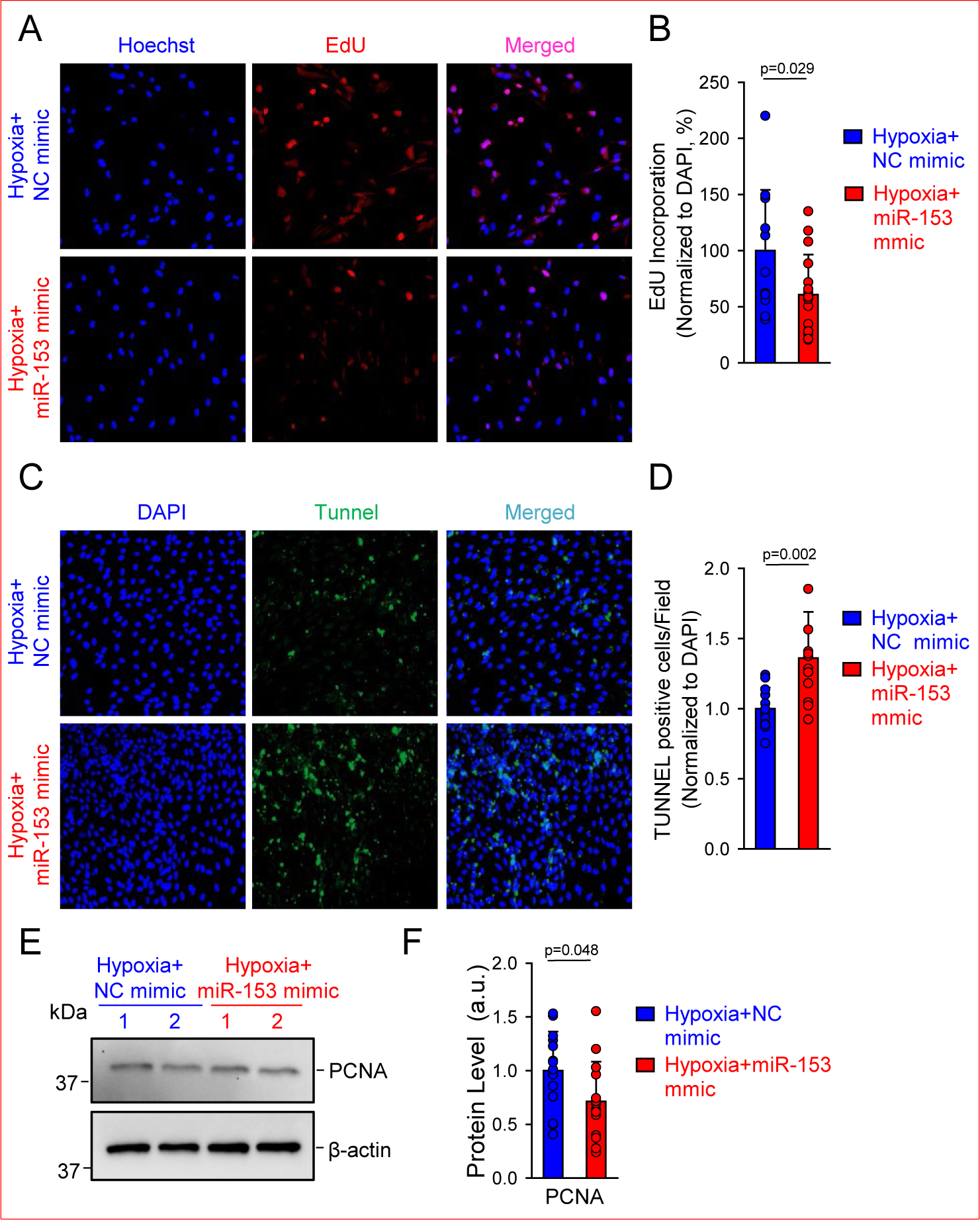
miR-153 reduces hypoxia-induced cell survival. **A-B:** Representative images (A) and summarized data (B) showing EdU incorporation in EC transfected with NC or miR-153 mimic and exposed to hypoxia for 72 hours (n=12-16 per group). **C-D:** Representative images (C) and summarized data (D) showing TUNEL staining in EC transfected with NC or miR-153 mimic and exposed to hypoxia for 72 hours (n=12 per group). **E-F:** Representative Western blot images (E) and summarized data (F) showing PCNA protein expression (normalized to β-actin) in EC transfected with NC or miR-153 mimic and exposed to hypoxia for 72 hours (n=14 per group). Values are mean±SD.

### miR-153 prevents EndMT and endothelial permeability induced by TGF-***β***1

TGF-β1 is well-known inducer of EndMT in different types of EC including lung vascular EC. We previously reported that TGF-β1 treatment for 7 days induced EndMT through upregulation of SNAI1/2, changed cell morphology and disrupted continuous adherens junctions in human lung vascular EC (19). Therefore, we quantified miR-153 level in EC treated with TGF-β1 for 7 days. Similar to hypoxic cells, TGF-β1-treated cells demonstrated the dramatically decreased expression of miR-153 (Fig. S1A). Furthermore, the transwell assay confirmed the promoting effect of TGF-β1 on endothelial permeability (Fig. S1B). In addition, we demonstrated that lung vascular EC isolated from patients with pulmonary arterial hypertension (PAH, Group 1 PH) (19) underwent EndMT program and exhibited the number of discontinuous adherens junctions similar to that we observed in TGF-β1–treated cells. Thus, we also compared miR-153 level in EC isolated from healthy subjects and PAH patients. Highly sensitive qPCR experiments revealed significantly lower miR-153 expression in PAH-EC compared with controls (0.44±0.19, p=0.028, Fig. S2) indicating the clinical relevance of the link between miR-153 and EndMT.

In this set of experiments, we aimed to examine the role of miR-153 in the development of EndMT. We applied loss- or gain-of-function approach to modulate level of miR-153 and then determined its effect on EndMT profile. As shown in Fig.6A and B, miR-153 inhibition resulted in the enhanced expression of SNAI1 (3.3-fold, *p*=<0.001), SNAI2 (3.4-fold, *p*=<0.001), and ACTA2 (2.2-fold, *p=*0.001), while reduced PECAM-1 expression (0.56-fold, *p*=<0.001). As expected, miR-153 inhibitor also negatively affected the endothelial barrier function in vehicle-treated cells which led to the enhanced permeability (Fig. 6C).

**Figure 6.**
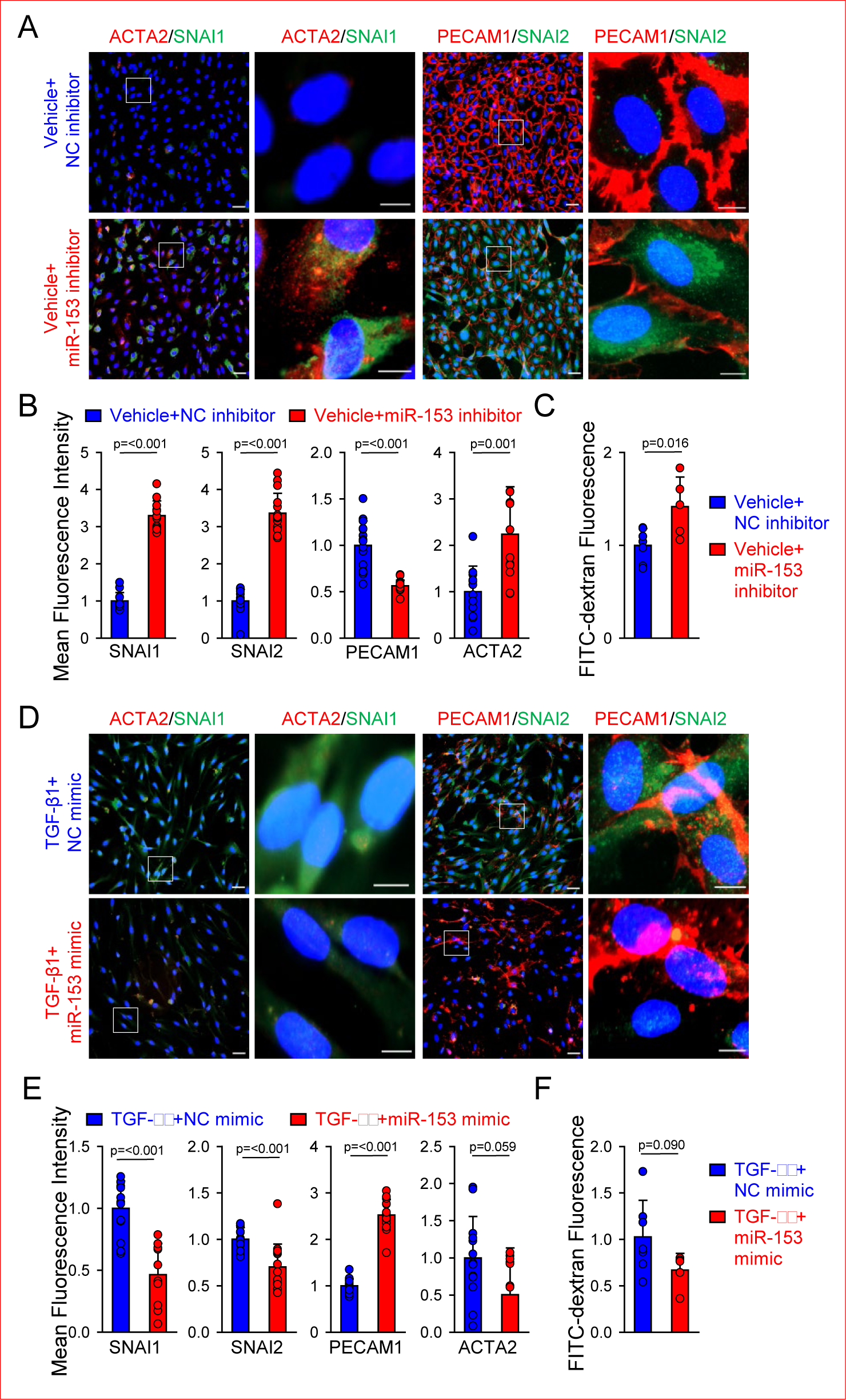
miR-153 abolishes TGF-β1-induced EndMT and permeability. **A-B:** Representative images (A) and summarized data (B) showing mean fluorescence intensity for SNAI1, SNAI2, PECAM1, and ACTA2 in human lung vascular EC transfected with NC or miR-153 inhibitor and treated with vehicle for 7 days (n=12-15 per group). Nuclei were counterstained with DAPI. Scale bars: 50 µm and 10 µm. **C:** Summarized data showing mean fluorescence intensity for FITC-dextran diffused through EC monolayer transfected with NC or miR-153 inhibitor and treated with vehicle for 7 days (n=5-7 per group). **D-E:** Representative images (A) and summarized data (B) showing mean fluorescence intensity for SNAI1, SNAI2, PECAM1, and ACTA2 in EC transfected with NC or miR-153 mimic and treated with TGF-β1 (10 ng/mL) for 7 days (n=10-15 per group). **F:** Summarized data showing mean fluorescence intensity for FITC-dextran diffused through the endothelial monolayer of EC transfected with NC or miR-153 mimic and treated with TGF-β1 (10 ng/mL) for 7 days (n=5-7 per group). Values are mean±SD.

Similar to EC exposed to hypoxia, miR-153 mimic significantly attenuated EndMT in cells exposed to TGF-β1. Indeed, miR-153 overexpression significantly suppressed TGF-β1-induced upregulation of SNAI1/SNAI2 and restored TGF-β1-induced downregulation of PECAM-1 (Fig. 6D and E). The level of ACTA2 showed the trend towards decreasing in miR-153-transfected cells compared with controls but did not reach significance (p=0.059). Similar to cell exposed to hypoxia, miR-153 had no significant effect on TGF-β1–induced endothelial permeability in EC (*p*=0.09), although a trend towards the restored barrier function was evident (Fig. 6F). Collectively, our findings demonstrate that miR-153 is crucial to maintain endothelial phenotype and barrier integrity in normal EC while miR-153 delivery may serve as a promising therapeutic strategy to attenuate EndMT and preserve endothelial identity in PH.

## Discussion

This study demonstrates the critical role of miR-153 in regulating EndMT and vascular permeability in human lung vascular endothelial cells (ECs). The principal findings are as follows: (i) decreased miR-153 levels are associated with hypoxia- or TGF-β1-induced EndMT; (ii) hypoxia or TGF-β1 disrupts cell-cell junctions, thereby compromising barrier integrity; (iii) loss of miR-153 induces spontaneous EndMT, increases endothelial permeability, and alters EC survival; (iv) miR-153 overexpression attenuates EndMT in response to hypoxia or TGF-β1; and (v) miR-153 expression is reduced in ECs isolated from patients with idiopathic PAH, a condition characterized by an enhanced EndMT profile.

TGF-β1 is widely recognized as a canonical inducer of both EMT and EndMT, contributing to the pathogenesis of numerous diseases, including cancer, fibrosis, systemic sclerosis, and PH (1, 3, 4). Although hypoxia is well known to promote EMT (22), its impact on EndMT has been less extensively explored. Previous studies have shown that hypoxic exposure (1–3% O) for up to seven days induces EndMT in rodent and human pulmonary vascular ECs in a time-dependent manner (23). Yu et al. reported concomitant SNAI1 upregulation and CDH5 downregulation in human pulmonary artery ECs as early as 6 hours after hypoxic exposure (24), further supporting evidence that SNAI1 directly binds to E-box promoter regions of cadherins (25, 26). Mesenchymal markers, including ACTA2, vimentin, and fibronectin, were significantly upregulated in human ECs after 12-24 hours of hypoxia (24). Similar findings were reported by Choi et al. (27), who concluded that hypoxia-induced EndMT develops within 24 hours. Collectively, these studies are consistent with our findings, though the specific outcomes depend on oxygen concentration, exposure duration, and vascular bed origin.

In our experiments, both TGF-β1- and hypoxia-induced EndMT not only altered the EC transcriptional profile but also impaired endothelial barrier integrity. Several studies have demonstrated that low oxygen levels or inflammatory cytokines increase paracellular permeability by disrupting tight and adherens junctions mediated by homophilic adhesion molecules such as occludin and claudin (28-30). Direct SNAI1-mediated repression of CDH5 weakens adherens junctions, resulting in increased vascular permeability. Chemical hypoxia (pseudohypoxia) induced by CoCl for 24-72 hours also promoted SNAI1-mediated EndMT, proliferation, and migration in human ECs (31, 32), although barrier function was not assessed. Similarly, hypoxia-induced EMT disrupts adherens junctions and enhances permeability in cancer cells (33, 34). Together, these findings suggest that disruption of endothelial cell-cell junctions represents an early trigger in EndMT initiation.

In vivo, the accumulation of α-SMA-positive, endothelial-derived cells within small pulmonary arterioles of hypoxia- or Sugen/hypoxia-induced PH models further supports the notion that EndMT-mediated junctional disassembly contributes to increased vascular permeability, inflammation, and vascular remodeling (24, 29, 35). Notably, EndMT susceptibility appears to be vessel-size dependent: small pulmonary arteries exhibit robust EndMT and smooth muscle-like cell emergence, whereas large pulmonary arteries show minimal changes, indicating regional heterogeneity in hypoxia-driven EC plasticity (36, 37). Lineage-tracing studies using Tie2-Cre;ROSA^tdTomato^ reporter mice confirmed that two weeks of hypoxic exposure drive endothelial transdifferentiation, with transitional EndMT cells accumulating in remodeled pulmonary arterioles (38). Collectively, these findings underscore hypoxia as a central driver of EC barrier dysfunction and pathological vascular remodeling, highlighting EndMT as a key mechanism linking oxygen deprivation to PH progression.

The functional consequences of oxygen deprivation on EC survival remain controversial, likely due to differences in EC subtype, oxygen level, and exposure duration. Zhang et al. reported a pro-proliferative response of macrovascular lung ECs to severe hypoxia (1% O) for 24 hours (39), whereas microvascular ECs underwent apoptosis under the same conditions. Another study demonstrated that microvascular lung ECs did not exhibit increased proliferation or migration after 48 hours of 1% O exposure (40). Interestingly, when microvascular ECs were pre-exposed to hypoxia for 7 days, followed by continued hypoxic stress for 48 hours, proliferative capacity increased significantly. In our study, we observed enhanced EC proliferation under physiologically relevant, moderate hypoxia (3% O) for 72 hours. This experimental design aligns with the transcriptional reprogramming and phenotypic changes characteristic of EndMT, while minimizing the cell loss observed under more severe hypoxia (1% O). Collectively, these findings suggest that short-term hypoxia (24-48 hours) induces EC injury and apoptosis, whereas prolonged hypoxia (3-7 days) promotes apoptosis-resistant ECs to adopt a mesenchymal phenotype with increased proliferation and migration.

Previous reports indicate that miR-153 functions primarily as a tumor suppressor and vascular regulator. MiR-153 inhibits proliferation, migration, invasion, and angiogenesis by targeting multiple oncogenic signaling pathways, including Jagged1, ADAM19, AKT, ROCK1, NFATc3, SNAI1, and ZEB2 (16, 41-45). Intriguingly, miR-153 has been shown to negatively regulate STAT3/VEGF signaling in tumor microenvironment EC by directly targeting 3’-UTR of VEGF-A and Cdc42, both critically involved in EC permeability and barrier function (46, 47). Using a dual-luciferase reporter assay, SNAI1 was also identified as a direct target of miR-153 (16, 48). Loss of miR-153 has been associated with enhanced epithelial-mesenchymal transition and metastasis, while its overexpression promotes apoptosis, reduces stem cell-like traits, and inhibits tumor growth both in vitro and in vivo. Moreover, reduced serum miR-153 levels have been correlated with poor clinical outcomes in cancer patients, suggesting its potential as a diagnostic and prognostic biomarker (16, 49).

In ECs and smooth muscle cells, downregulation of miR-153 facilitates hypoxia-induced angiogenesis and vascular remodeling, underscoring its role as a brake on maladaptive vascular responses (41, 42, 50). Our findings are consistent with these studies, demonstrating that miR-153 preserves lung vascular EC function by inhibiting EndMT and reducing endothelial permeability in response to hypoxia or inflammatory cytokines. Importantly, VEGF-associated pulmonary vascular leakage has been observed in lung tissues from PH patients (35). In the present study, we found that miR-153 expression was significantly downregulated in lung vascular ECs isolated from PH patients. Loss of miR-153 may impair the regenerative capacity of ECs and enhance their interaction with smooth muscle cells and fibroblasts, thereby amplifying vascular remodeling in PH.

## Conclusions

Our findings, for the first time, identify miR-153 as a pivotal mechanistic link between EndMT, EC dysfunction, and pulmonary vascular pathology. Collectively, the data support the concept that miR-153 functions as an endothelial gatekeeper and may serve as both a prognostic biomarker and a promising therapeutic target for the treatment of PH.

## ACKNOWLEDGMENTS

This study was supported partially by grants from the University of Minnesota Hormel Institute Eagles Telethon Postdoctoral Fellowship (I.E.), University of Minnesota Hormel Institute Transformative Ideas Program (A.B.), American Heart Association Undergraduate Student Training Award 25IAUST1363304 (A.B.), Minnesota Partnership for Biotechnology and Medical Genomics program MNP#24.04 (A.B. and C.M.P), the National Lung, Heart, and Blood Institute of the National Institutes of Health HL135807 (J.X.-J.Y.).

---

**TABLE S1.**
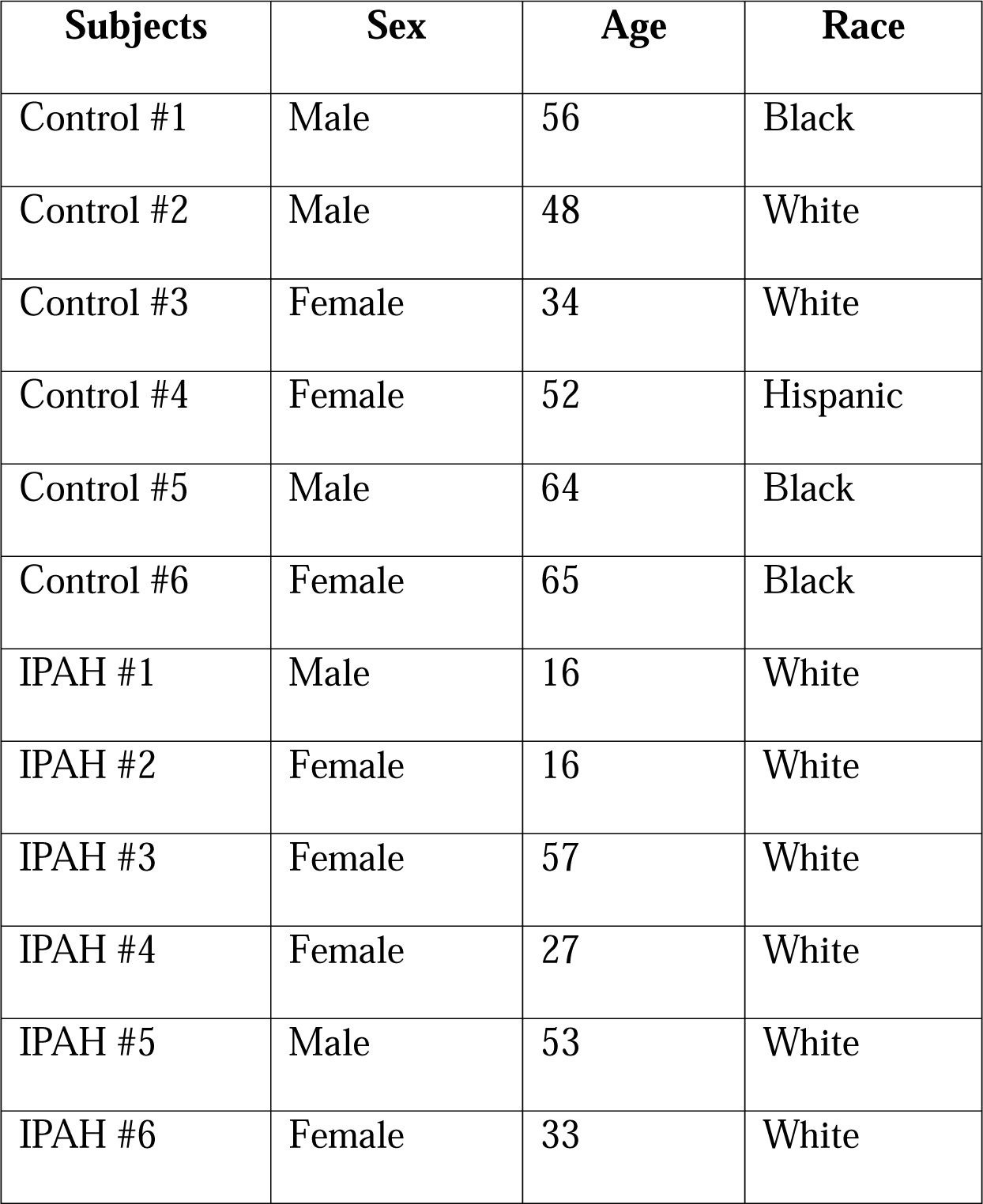
Demographic information of human donors.

## FIGURE LEGENDS

**Figure S1.**
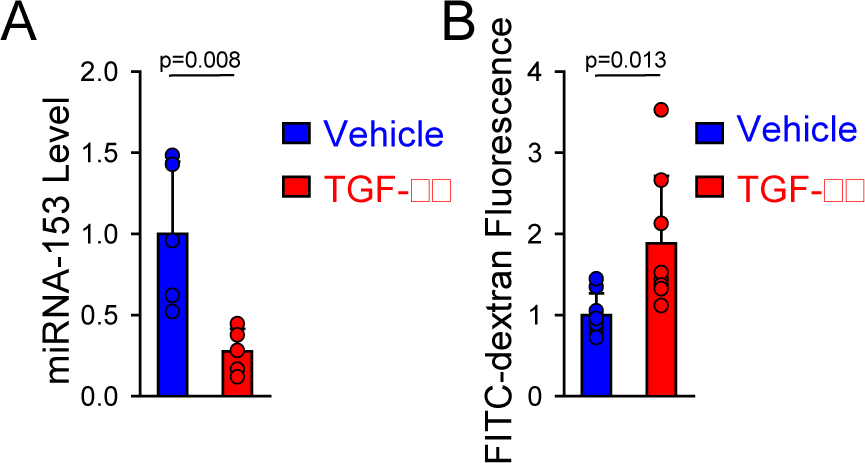
**A:** Summarized data showing RT-PCR analysis of miR-153 expression (normalized to U6) in EC treated with vehicle or TGF-β1 (10 ng/mL) for 7 days (n=5 per group). **B:** Summarized data showing mean fluorescence intensity for FITC-dextran diffused through the endothelial monolayer of EC treated with vehicle or TGF-β1 for 7 days (n=8 per group). Values are mean±SD.

**Figure S2.**
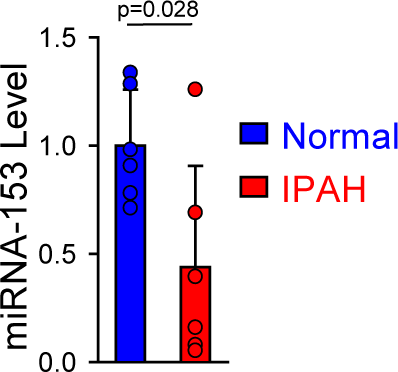
Summarized data showing RT-PCR analysis of miR-153 (normalized to U6) in lung vascular EC isolated from control subjects and idiopathic PAH patients (n=6 per group). Values are mean±SD.

## References

1 Kovacic JC, Dimmeler S, Harvey RP, Finkel T, Aikawa E, Krenning G, Baker AH. Endothelial to Mesenchymal Transition in Cardiovascular Disease: JACC State-of-the-Art Review. J Am Coll Cardiol 2019; 73: 190–209.

2 Giordanengo L, Proment A, Botta V, Picca F, Munir HMW, Tao J, Olivero M, Taulli R, Bersani F, Sangiolo D, Novello S, Scagliotti GV, Merlini A, Doronzo G. Shifting Shapes: The Endothelial-to-Mesenchymal Transition as a Driver for Cancer Progression. Int J Mol Sci 2025; 26.

3 Gorelova A, Berman M, Al Ghouleh I. Endothelial-to-Mesenchymal Transition in Pulmonary Arterial Hypertension. Antioxid Redox Signal 2021; 34: 891–914.

4 Li Y, Lui KO, Zhou B. Reassessing endothelial-to-mesenchymal transition in cardiovascular diseases. Nat Rev Cardiol 2018; 15: 445–456.

5 Hulshoff MS, Xu X, Krenning G, Zeisberg EM. Epigenetic Regulation of Endothelial-to-Mesenchymal Transition in Chronic Heart Disease. Arterioscler Thromb Vasc Biol 2018; 38: 1986–1996.

6 Zhang H, Hu J, Liu L. MiR-200a modulates TGF-beta1-induced endothelial-to-mesenchymal shift via suppression of GRB2 in HAECs. Biomed Pharmacother 2017; 95: 215–222.

7 Zhang J, Zeng Y, Chen J, Cai D, Chen C, Zhang S, Chen Z. miR-29a/b cluster suppresses high glucose-induced endothelial-mesenchymal transition in human retinal microvascular endothelial cells by targeting Notch2. Exp Ther Med 2019; 17: 3108–3116.

8 Jordan NP, Tingle SJ, Shuttleworth VG, Cooke K, Redgrave RE, Singh E, Glover EK, Ahmad Tajuddin HB, Kirby JA, Arthur HM, Ward C, Sheerin NS, Ali S. MiR-126-3p Is Dynamically Regulated in Endothelial-to-Mesenchymal Transition during Fibrosis. Int J Mol Sci 2021; 22.

9 Wang Y, Sun L, Jia X, Yang M, Xu W. miR-29 inhibits endothelial-to-mesenchymal transition in pulmonary hypertension of the newborn by regulating LRP6. FASEB J 2025; 39: e70502.

10 Xu YP, He Q, Shen Z, Shu XL, Wang CH, Zhu JJ, Shi LP, Du LZ. MiR-126a-5p is involved in the hypoxia-induced endothelial-to-mesenchymal transition of neonatal pulmonary hypertension. Hypertens Res 2017; 40: 552–561.

11 Tang H, Mao J, Ye X, Zhang F, Kerr WG, Zheng T, Zhu Z. SHIP-1, a target of miR-155, regulates endothelial cell responses in lung fibrosis. FASEB J 2020; 34: 2011–2023.

12 Yousefnia S. A comprehensive review on miR-153: Mechanistic and controversial roles of miR-153 in tumorigenicity of cancer cells. Front Oncol 2022; 12: 985897.

13 Zhang Z, Sun J, Bai Z, Li H, He S, Chen R, Che X. MicroRNA-153 acts as a prognostic marker in gastric cancer and its role in cell migration and invasion. Onco Targets Ther 2015; 8: 357–364.

14 Li W, Zhai L, Zhao C, Lv S. MiR-153 inhibits epithelial-mesenchymal transition by targeting metadherin in human breast cancer. Breast Cancer Res Treat 2015; 150: 501–509.

15 Xia W, Ma X, Li X, Dong H, Yi J, Zeng W, Yang Z. miR-153 inhibits epithelial-to-mesenchymal transition in hepatocellular carcinoma by targeting Snail. Oncol Rep 2015; 34: 655–662.

16 Xu Q, Sun Q, Zhang J, Yu J, Chen W, Zhang Z. Downregulation of miR-153 contributes to epithelial-mesenchymal transition and tumor metastasis in human epithelial cancer. Carcinogenesis 2013; 34: 539–549.

17 Babicheva A, Elmadbouh I, Song S, Thompson MA, Powers R, Jain PP, Izadi A, Chen J, Yung L, Parmisano S, Paquin C, Wang WT, Chen Y, Wang T, Alotaibi M, Shyy JY, Thistlethwaite PA, Wang J, Makino A, Prakash YS, Pabelick CM, Yuan JX. Store-operated Ca(2+) entry is involved in endothelium-to-mesenchymal transition in lung vascular endothelial cells. Am J Physiol Lung Cell Mol Physiol 2025; 328: L844–L857.

18 Babicheva A, Ayon RJ, Zhao T, Ek Vitorin JF, Pohl NM, Yamamura A, Yamamura H, Quinton BA, Ba M, Wu L, Ravellette KS, Rahimi S, Balistrieri F, Harrington A, Vanderpool RR, Thistlethwaite PA, Makino A, Yuan JX. MicroRNA-mediated downregulation of K(+) channels in pulmonary arterial hypertension. Am J Physiol Lung Cell Mol Physiol 2020; 318: L10–L26.

19 Tang H, Babicheva A, McDermott KM, Gu Y, Ayon RJ, Song S, Wang Z, Gupta A, Zhou T, Sun X, Dash S, Wang Z, Balistrieri A, Zheng Q, Cordery AG, Desai AA, Rischard F, Khalpey Z, Wang J, Black SM, Garcia JGN, Makino A, Yuan JX. Endothelial HIF-2alpha contributes to severe pulmonary hypertension due to endothelial-to-mesenchymal transition. Am J Physiol Lung Cell Mol Physiol 2018; 314: L256–L275.

20 Song S, Carr SG, McDermott KM, Rodriguez M, Babicheva A, Balistrieri A, Ayon RJ, Wang J, Makino A, Yuan JX. STIM2 (Stromal Interaction Molecule 2)-Mediated Increase in Resting Cytosolic Free Ca(2+) Concentration Stimulates PASMC Proliferation in Pulmonary Arterial Hypertension. Hypertension 2018; 71: 518–529.

21 Wang L, Astone M, Alam SK, Zhu Z, Pei W, Frank DA, Burgess SM, Hoeppner LH. Suppressing STAT3 activity protects the endothelial barrier from VEGF-mediated vascular permeability. Dis Model Mech 2021; 14.

22 Hapke RY, Haake SM. Hypoxia-induced epithelial to mesenchymal transition in cancer. Cancer Lett 2020; 487: 10–20.

23 Evrard SM, Lecce L, Michelis KC, Nomura-Kitabayashi A, Pandey G, Purushothaman KR, d’Escamard V, Li JR, Hadri L, Fujitani K, Moreno PR, Benard L, Rimmele P, Cohain A, Mecham B, Randolph GJ, Nabel EG, Hajjar R, Fuster V, Boehm M, Kovacic JC. Endothelial to mesenchymal transition is common in atherosclerotic lesions and is associated with plaque instability. Nat Commun 2016; 7: 11853.

24 Yu M, Peng L, Liu P, Yang M, Zhou H, Ding Y, Wang J, Huang W, Tan Q, Wang Y, Xie W, Kong H, Wang H. Paeoniflorin Ameliorates Chronic Hypoxia/SU5416-Induced Pulmonary Arterial Hypertension by Inhibiting Endothelial-to-Mesenchymal Transition. Drug Des Devel Ther 2020; 14: 1191–1202.

25 Cano A, Perez-Moreno MA, Rodrigo I, Locascio A, Blanco MJ, del Barrio MG, Portillo F, Nieto MA. The transcription factor snail controls epithelial-mesenchymal transitions by repressing E-cadherin expression. Nat Cell Biol 2000; 2: 76–83.

26 Lin T, Ponn A, Hu X, Law BK, Lu J. Requirement of the histone demethylase LSD1 in Snai1-mediated transcriptional repression during epithelial-mesenchymal transition. Oncogene 2010; 29: 4896–4904.

27 Choi SH, Hong ZY, Nam JK, Lee HJ, Jang J, Yoo RJ, Lee YJ, Lee CY, Kim KH, Park S, Ji YH, Lee YS, Cho J, Lee YJ. A Hypoxia-Induced Vascular Endothelial-to-Mesenchymal Transition in Development of Radiation-Induced Pulmonary Fibrosis. Clin Cancer Res 2015; 21: 3716–3726.

28 Sun X, Nakajima E, Norbrun C, Sorkhdini P, Yang AX, Yang D, Ventetuolo CE, Braza J, Vang A, Aliotta J, Banerjee D, Pereira M, Baird G, Lu Q, Harrington EO, Rounds S, Lee CG, Yao H, Choudhary G, Klinger JR, Zhou Y. Chitinase 3 like 1 contributes to the development of pulmonary vascular remodeling in pulmonary hypertension. JCI Insight 2022; 7.

29 Mammoto T, Muyleart M, Konduri GG, Mammoto A. Twist1 in Hypoxia-induced Pulmonary Hypertension through Transforming Growth Factor-beta-Smad Signaling. Am J Respir Cell Mol Biol 2018; 58: 194–207.

30 Good RB, Gilbane AJ, Trinder SL, Denton CP, Coghlan G, Abraham DJ, Holmes AM. Endothelial to Mesenchymal Transition Contributes to Endothelial Dysfunction in Pulmonary Arterial Hypertension. Am J Pathol 2015; 185: 1850–1858.

31 Zou R, Feng YF, Xu YH, Shen MQ, Zhang X, Yuan YZ. Yes-associated protein promotes endothelial-to-mesenchymal transition of endothelial cells in choroidal neovascularization fibrosis. Int J Ophthalmol 2022; 15: 701–710.

32 Song S, Ji Y, Zhang G, Zhang X, Li B, Li D, Jiang W. Protective Effect of Atazanavir Sulphate Against Pulmonary Fibrosis In Vivo and In Vitro. Basic Clin Pharmacol Toxicol 2018; 122: 199–207.

33 Medici D, Kalluri R. Endothelial-mesenchymal transition and its contribution to the emergence of stem cell phenotype. Semin Cancer Biol 2012; 22: 379–384.

34 Ikenouchi J, Matsuda M, Furuse M, Tsukita S. Regulation of tight junctions during the epithelium-mesenchyme transition: direct repression of the gene expression of claudins/occludin by Snail. J Cell Sci 2003; 116: 1959–1967.

35 Zhou W, Liu K, Zeng L, He J, Gao X, Gu X, Chen X, Jing Li J, Wang M, Wu D, Cai Z, Claesson-Welsh L, Ju R, Wang J, Zhang F, Chen Y. Targeting VEGF-A/VEGFR2 Y949 Signaling-Mediated Vascular Permeability Alleviates Hypoxic Pulmonary Hypertension. Circulation 2022; 146: 1855–1881.

36 Wang EL, Zhang JJ, Luo FM, Fu MY, Li D, Peng J, Liu B. Cerebellin-2 promotes endothelial-mesenchymal transition in hypoxic pulmonary hypertension rats by activating NF-kappaB/HIF-1alpha/Twist1 pathway. Life Sci 2023; 328: 121879.

37 Huang J, Lu W, Ouyang H, Chen Y, Zhang C, Luo X, Li M, Shu J, Zheng Q, Chen H, Chen J, Tang H, Sun D, Yuan JX, Yang K, Wang J. Transplantation of Mesenchymal Stem Cells Attenuates Pulmonary Hypertension by Normalizing the Endothelial-to-Mesenchymal Transition. Am J Respir Cell Mol Biol 2020; 62: 49–60.

38 Woo KV, Shen IY, Weinheimer CJ, Kovacs A, Nigro J, Lin CY, Chakinala M, Byers DE, Ornitz DM. Endothelial FGF signaling is protective in hypoxia-induced pulmonary hypertension. J Clin Invest 2021; 131.

39 Zhang Q, Yaoita N, Tabuchi A, Liu S, Chen SH, Li Q, Hegemann N, Li C, Rodor J, Timm S, Laban H, Finkel T, Stevens T, Alvarez DF, Erfinanda L, de Perrot M, Kucherenko MM, Knosalla C, Ochs M, Dimmeler S, Korff T, Verma S, Baker AH, Kuebler WM. Endothelial Heterogeneity in the Response to Autophagy Drives Small Vessel Muscularization in Pulmonary Hypertension. Circulation 2024; 150: 466–487.

40 Zhang B, Niu W, Dong HY, Liu ML, Luo Y, Li ZC. Hypoxia induces endothelial mesenchymal transition in pulmonary vascular remodeling. Int J Mol Med 2018; 42: 270–278.

41 Lu Y, Li D, Shan L. MicroRNA153 induces apoptosis by targeting NFATc3 to improve vascular remodeling in pulmonary hypertension. Clin Exp Hypertens 2023; 45: 2140810.

42 Zhao M, Wang W, Lu Y, Wang N, Kong D, Shan L. MicroRNA 153 attenuates hypoxia induced excessive proliferation and migration of pulmonary arterial smooth muscle cells by targeting ROCK1 and NFATc3. Mol Med Rep 2021; 23.

43 Liang H, Ge F, Xu Y, Xiao J, Zhou Z, Liu R, Chen C. miR-153 inhibits the migration and the tube formation of endothelial cells by blocking the paracrine of angiopoietin 1 in breast cancer cells. Angiogenesis 2018; 21: 849–860.

44 Shan N, Shen L, Wang J, He D, Duan C. MiR-153 inhibits migration and invasion of human non-small-cell lung cancer by targeting ADAM19. Biochem Biophys Res Commun 2015; 456: 385–391.

45 Yuan Y, Du W, Wang Y, Xu C, Wang J, Zhang Y, Wang H, Ju J, Zhao L, Wang Z, Lu Y, Cai B, Pan Z. Suppression of AKT expression by miR-153 produced anti-tumor activity in lung cancer. Int J Cancer 2015; 136: 1333–1340.

46 Zhang W, Mao S, Shi D, Zhang J, Zhang Z, Guo Y, Wu Y, Wang R, Wang L, Huang Y, Yao X. MicroRNA-153 Decreases Tryptophan Catabolism and Inhibits Angiogenesis in Bladder Cancer by Targeting Indoleamine 2,3-Dioxygenase 1. Front Oncol 2019; 9: 619.

47 Ma Y, Xue Y, Liu X, Qu C, Cai H, Wang P, Li Z, Li Z, Liu Y. SNHG15 affects the growth of glioma microvascular endothelial cells by negatively regulating miR-153. Oncol Rep 2017; 38: 3265–3277.

48 Zhang B, Fu T, Zhang L. MicroRNA-153 suppresses human laryngeal squamous cell carcinoma migration and invasion by targeting the SNAI1 gene. Oncol Lett 2018; 16: 5075–5083.

49 Zhao G, Zhang Y, Zhao Z, Cai H, Zhao X, Yang T, Chen W, Yao C, Wang Z, Wang Z, Han C, Wang H. MiR-153 reduces stem cell-like phenotype and tumor growth of lung adenocarcinoma by targeting Jagged1. Stem Cell Res Ther 2020; 11: 170.

50 Li L, Wang M, Mei Z, Cao W, Yang Y, Wang Y, Wen A. lncRNAs HIF1A-AS2 facilitates the up-regulation of HIF-1alpha by sponging to miR-153-3p, whereby promoting angiogenesis in HUVECs in hypoxia. Biomed Pharmacother 2017; 96: 165–172.

