## Supplemental Tables 2 and 3 for "Loss of miRNA-153 promotes endothelial-to-mesenchymal transition and compromises lung vascular integrity"

**TABLE S2.** *In silico* analysis to predict miRNAs that directly targets 3’-UTR of human SNAI1 gene.

| **Target miRNA** | **Position of**  **3'-UTR** | **Complementary sequence of target region (top) and miRNA (bottom)** | **Context++ score** | **Context++ score percentile** |
| --- | --- | --- | --- | --- |
| hsa-miR-153-3p | 440-447 | 5'    ...ACGAGGUGUGACUAACUAUGCAA... | -0.49 | 99 |
|  |  | \|\|\|\|\|\|\| |  |  |
|  |  | 3'        CUAGUGAAAACACUGAUACGUU |  |  |
| hsa-miR-22-3p | 522-529 | 5'    ...GAUGCCCCGAGCCCAGGCAGCUA... | -0.45 | 98 |
|  |  | \|\|\|\|\|\|\| |  |  |
|  |  | 3'        UGUCAAGAAGUUGACCGUCGAA |  |  |
| hsa-miR-30b-5p | 706-713 | 5' ...GGCCUGGGAGGAAGAUGUUUACA... | -0.4 | 98 |
|  |  | \|\|\|\|\|\|\| |  |  |
|  |  | 3'     GAAGGUCAGCCCCUACAAAUGU |  |  |
| hsa-miR-30d-5p | 706-713 | 5'   ...GGCCUGGGAGGAAGAUGUUUACA... | -0.39 | 98 |
|  |  | \|\|\|\|\|\|\| |  |  |
|  |  | 3'      CGACUCUCACAUCCUACAAAUGU |  |  |
| hsa-miR-30c-5p | 706-713 | 5'   ...GGCCUGGGAGGAAGAUGUUUACA... | -0.4 | 98 |
|  |  | \|\|\|\|\|\|\| |  |  |
|  |  | 3'       GAAGGUCAGCUCCUACAAAUGU |  |  |
| hsa-miR-30e-5p | 706-713 | 5'   ...GGCCUGGGAGGAAGAUGUUUACA... | -0.4 | 98 |
|  |  | \|\|\|\|\|\|\| |  |  |
|  |  | 3'       UCGACUCACAUCCUACAAAUGU |  |  |
| hsa-miR-30a-5p | 706-713 | 5'  ...GGCCUGGGAGGAAGAUGUUUACA... | -0.41 | 98 |
|  |  | \|\|\|\|\|\|\| |  |  |
|  |  | 3'      GAAGGUCAGUUCCUACAAAUGU |  |  |
| hsa-miR-199b-5p | 725-731 | 5'  ...UACAUUUUUAAAGGUACACUGGU... | -0.42 | 97 |
|  |  | \|\|\|\|\|\|\| |  |  |
|  |  | 3'     CUUGUCUAUCAGAUUUGUGACCC |  |  |
| hsa-miR-199a-5p | 725-731 | 5'  ...UACAUUUUUAAAGGUACACUGGU... | -0.42 | 97 |
|  |  | \|\|\|\|\|\|\| |  |  |
|  |  | 3'     CUUGUCCAUCAGACUUGUGACCC |  |  |
| hsa-miR-27b-3p | 595-601 | 5'   ...AAUGUCUGAAAAGGGACUGUGAG... | -0.16 | 89 |
|  |  | \|\|\|\|\|\|\| |  |  |
|  |  | 3'        CGUCUUGAAUCGGUGACACUU |  |  |
| hsa-miR-27a-3p | 595-601 | 5'   ...AAUGUCUGAAAAGGGACUGUGAG... | -0.16 | 89 |
|  |  | \|\|\|\|\|\|\| |  |  |
|  |  | 3'        CGCCUUGAAUCGGUGACACUU |  |  |
| hsa-miR-363-3p | 443-449 | 5'  ...AGGUGUGACUAACUAUGCAAUAA... | -0.16 | 88 |
|  |  | \|\|\|\|\|\| |  |  |
|  |  | 3'     AUGUCUACCUAUGGCACGUUAA |  |  |
| hsa-miR-367-3p | 443-449 | 5'  ...AGGUGUGACUAACUAUGCAAUAA... | -0.17 | 88 |
|  |  | \|\|\|\|\|\| |  |  |
|  |  | 3'     AGUGGUAACGAUUUCACGUUAA |  |  |
| hsa-miR-25-3p | 443-449 | 5'  ...AGGUGUGACUAACUAUGCAAUAA... | -0.15 | 88 |
|  |  | \|\|\|\|\|\| |  |  |
|  |  | 3'     AGUCUGGCUCUGUUCACGUUAC |  |  |
| hsa-miR-32-3p | 443-449 | 5'   ...AGGUGUGACUAACUAUGCAAUAA... | -0.13 | 86 |
|  |  | \|\|\|\|\|\| |  |  |
|  |  | 3'      UGUCCGGCCCUGUUCACGUUAU |  |  |
| hsa-miR-92a-5p | 443-449 | 5'   ...AGGUGUGACUAACUAUGCAAUAA... | -0.12 | 86 |
|  |  | \|\|\|\|\|\| |  |  |
|  |  | 3'      ACGUUGAAUCAUUACACGUUAU |  |  |
| hsa-miR-92b-3p | 443-449 | 5'   ...AGGUGUGACUAACUAUGCAAUAA... | -0.12 | 86 |
|  |  | \|\|\|\|\|\| |  |  |
|  |  | 3'      CCUCCGGCCCUGCUCACGUUAU |  |  |
| hsa-miR-3681-3p | 595-601 | 5' ...AAUGUCUGAAAAGGGACUGUGAG... | -0.15 | 86 |
|  |  | \|\|\|\|\|\| |  |  |
|  |  | 3'    UCAUCACCUACUUCGUGACACA |  |  |
| hsa-miR-128-3p | 595-601 | 5'  ...AAUGUCUGAAAAGGGACUGUGAG... | -0.12 | 83 |
|  |  | \|\|\|\|\|\| |  |  |
|  |  | 3'      UUUCUCUGGCCAAGUGACACU |  |  |
| hsa-miR-216a-3p | 595-601 | 5'   ...AAUGUCUGAAAAGGGACUGUGAG... | -0.11 | 81 |
|  |  | \|\|\|\|\|\| |  |  |
|  |  | 3'      UAUUAGGGUCUCUGGUGACACU |  |  |

**TABLE S3.** *In silico* analysis to predict miRNAs that directly targets 3’-UTR of human SNAI2 gene.

| **Target gene** | **Position of 3' UTR** | **Complementary sequence of target region (top) and miRNA (bottom)** | **Context++ score** | **Context++ score percentile** |
| --- | --- | --- | --- | --- |
| hsa-miR-203a-3p.2 | 351-358 | 5' ...ACAUUGCUGCCAAAUCAUUUCAA... | -0.36 | 99 |
|  |  | \|\|\|\|\|\|\| |  |  |
|  |  | 3'     GAUCACCAGGAUUUGUAAAGUG |  |  |
| hsa-miR-124-3p.1 | 439-446 | 5'    ...CAGACGCGAACUCAGGUGCCUUA... | -0.48 | 99 |
|  |  | \|\|\|\|\|\|\| |  |  |
|  |  | 3'          CCGUAAGUGGCGCACGGAAU |  |  |
| hsa-miR-124-3p.1 | 843-849 | 5' ...CAAUACCCCUUCCAA--GUGCCUUU... | -0.48 | 99 |
|  |  | \|\|\|     \|\|\|\|\|\|\| |  |  |
|  |  | 3'         CCGUAAGUGGCGCACGGAAU |  |  |
| hsa-miR-124-3p.1 | 439-446 | 5'    ...CAGACGCGAACUCAGGUGCCUUA... | -0.48 | 99 |
|  |  | \|\|\|\|\|\|\| |  |  |
|  |  | 3'          CCGUAAGUGGCGCACGGAAU |  |  |
| hsa-miR-124-3p.1 | 843-849 | 5' ...CAAUACCCCUUCCAA--GUGCCUUU... | -0.48 | 99 |
|  |  | \|\|\|     \|\|\|\|\|\|\| |  |  |
|  |  | 3'         CCGUAAGUGGCGCACGGAAU |  |  |
| hsa-miR-124-3p.1 | 439-446 | 5'    ...CAGACGCGAACUCAGGUGCCUUA... | -0.48 | 99 |
|  |  | \|\|\|\|\|\|\| |  |  |
|  |  | 3'          CCGUAAGUGGCGCACGGAAU |  |  |
| hsa-miR-124-3p.1 | 843-849 | 5' ...CAAUACCCCUUCCAA--GUGCCUUU... | -0.48 | 99 |
|  |  | \|\|\|     \|\|\|\|\|\|\| |  |  |
|  |  | 3'         CCGUAAGUGGCGCACGGAAU |  |  |
| hsa-miR-33a-5p | 816-822 | 5'  ...CAAAUGCAUACCACAAAUGCAAU... | -0.2 | 97 |
|  |  | \|\|\|\|\|\|       \|\|\|\|\|\| |  |  |
|  |  | 3'    ACGUUACGUUGAUG--UUACGUG |  |  |
| hsa-miR-33b-5p | 816-822 | 5' ...CAAAUGCAUACCACAAAUGCAAU... | -0.2 | 97 |
|  |  | \|\|\|\|\|\|       \|\|\|\|\|\| |  |  |
|  |  | 3'    CGUUACGUUGUCG--UUACGUG |  |  |
| hsa-miR-124-3p.1 | 639-645 | 5' ...GCCAAUUAACAGUAUGUGCCUUG... | -0.32 | 96 |
|  |  | \|\|\|\|\|\|\| |  |  |
|  |  | 3'       CCGUAAGUGGCGCACGGAAU |  |  |
| hsa-miR-124-3p.1 | 639-645 | 5' ...GCCAAUUAACAGUAUGUGCCUUG... | -0.32 | 96 |
|  |  | \|\|\|\|\|\|\| |  |  |
|  |  | 3'       CCGUAAGUGGCGCACGGAAU |  |  |
| hsa-miR-124-3p.1 | 639-645 | 5' ...GCCAAUUAACAGUAUGUGCCUUG... | -0.32 | 96 |
|  |  | \|\|\|\|\|\|\| |  |  |
|  |  | 3'       CCGUAAGUGGCGCACGGAAU |  |  |
| hsa-miR-181c-5p | 587-593 | 5' ...GAACAAAACACAGGA-GAAUGUAU... | -0.26 | 96 |
|  |  | \|\|\|\|\|  \|\|\|\|\|\| |  |  |
|  |  | 3'     UGAGUGGCUGUCCAACUUACAA |  |  |
| hsa-miR-181b-5p | 587-593 | 5' ...GAACAAAACACAGGA--GAAUGUAU... | -0.26 | 96 |
|  |  | \|\|\|\|    \|\|\|\|\|\| |  |  |
|  |  | 3'     UGGGUGGCUGUCGUUACUUACAA |  |  |
| hsa-miR-181a-5p | 587-593 | 5' ...GAACAAAACACAGGA--GAAUGUAU... | -0.26 | 96 |
|  |  | \|\|\|\|    \|\|\|\|\|\| |  |  |
|  |  | 3'     UGAGUGGCUGUCGCAACUUACAA |  |  |
| hsa-miR-181d-5p | 587-593 | 5'  ...GAACAAAACACAGGAGAAUGUAU... | -0.25 | 95 |
|  |  | \|\|\|     \|\|\|\|\|\| |  |  |
|  |  | 3'    UGGGUGGCUGUUGUUACUUACAA |  |  |
| hsa-miR-4262 | 587-593 | 5'      ...GAACAAAACACAGGAGAAUGUAU... | -0.17 | 95 |
|  |  | \|\|\|\|\|\| |  |  |
|  |  | 3'              GUCCAUCAGACUUACAG |  |  |
| hsa-miR-206 | 673-679 | 5' ...AAAGAUUAGCUUUGA-ACAUUCCU... | -0.27 | 94 |
|  |  | \|\|\|    \|\|\|\|\|\|\| |  |  |
|  |  | 3'      GGUGUGUGAAGGAAUGUAAGGU |  |  |
| hsa-miR-1-3p | 673-679 | 5' ...AAAGAUUAGCUUUGA-ACAUUCCU... | -0.27 | 94 |
|  |  | \|\|\|    \|\|\|\|\|\|\| |  |  |
|  |  | 3'      UAUGUAUGAAGAAAUGUAAGGU |  |  |
| hsa-miR-200b-3p | 369-375 | 5' ...UUCAACUGAAAAGAACAGUAUUG... | -0.17 | 93 |
|  |  | \|\|\|\|\|\|\| |  |  |
|  |  | 3'     AGUAGUAAUGGUCCGUCAUAAU |  |  |
| hsa-miR-429 | 369-375 | 5' ...UUCAACUGAAAAGAACAGUAUUG... | -0.17 | 93 |
|  |  | \|\|\|\|\|\|\| |  |  |
|  |  | 3'     UGCCAAAAUGGUCUGUCAUAAU |  |  |
| hsa-miR-613 | 673-679 | 5'    ...AAAGAUUAGCUUUGAACAUUCCU... | -0.25 | 93 |
|  |  | \|\|\|\|\|\|\| |  |  |
|  |  | 3'          CCGUUUCUUCCUUGUAAGGA |  |  |
| hsa-miR-200c-3p | 369-375 | 5' ...UUCAACUGAAAAGAACAGUAUUG... | -0.16 | 92 |
|  |  | \|\|\|\|\|\|\| |  |  |
|  |  | 3'    AGGUAGUAAUGGGCCGUCAUAAU |  |  |
| hsa-miR-203a-3p.1 | 816-822 | 5' ...ACAUUGCUGCCAAAUCAUUUCAA... | -0.1 | 86 |
|  |  | \|\|\|\|\|\| |  |  |
|  |  | 3'     GAUCACCAGGAUUUGUAAAGU |  |  |
